# Nutrient and host availability drive the evolution of viral plasticity

**DOI:** 10.64898/2026.11.24.768896

**Authors:** Julie D. Pourtois, Tapan Goel, Joshua S. Weitz

## Abstract

Viruses of microbes inject their genome into host cells, take over host metabolism, and produce progeny virions that are released into the environment upon lysis. Environmental and host conditions shape viral traits, including the intracellular maturation rate of virions and the time between infection and lysis. The dependence of viral traits on environmental context, i.e., phenotypic plasticity, would appear to be inevitable for obligate intracellular parasites, which depend on host metabolic activity for replication. However, viruses, including viruses of microbes, encode genes that can decouple their replication from the host metabolic state. Here, we present a trait analysis and modeling framework to investigate how plasticity in maturation and lysis rates influences viral growth, invasion, and coexistence. First, analysis of experimental viral trait data shows support for high maturation rate plasticity and intermediate lysis rate plasticity across virus-microbe pairs. Next, leveraging an eco-evolutionary model, we find that viruses with high maturation rate plasticity outcompete other viruses when nutrient inflow is high or variable but not when nutrient inflow is low and stable. In addition, we find that high lysis rate plasticity maximizes viral invasion fitness and growth when nutrient density is high and susceptible cell density is low, or when environmental conditions are highly variable. These findings suggest that while plasticity is evolutionarily advantageous in a broad range of environments, decoupling viral traits (including both maturation and lysis rates) from host metabolism can be advantageous, even if costly, in environments with low resource inflow. Finally, we find that variation in plasticity can facilitate coexistence between viral types given a shared, susceptible host population. Overall, these results highlight the adaptive value of (de)coupling viral traits from environmental and host state and suggest new directions for exploring the link between plasticity and diversity for virus-microbe communities.

## Introduction

Viruses of microbes are obligate intracellular parasites. Their life history traits depend on the metabolic machinery of the host, including ribosomes and ATP levels, to replicate the viral genome and produce the proteins required for virion self-assembly and lysis. This metabolic dependence often results in a direct relationship between the rate of viral production inside the cell (the maturation rate), and the growth rate of the host cell, which itself varies with nutrient conditions [1, 2, 3]. Changes in the viral maturation rate in nutrient-poor conditions typically decrease burst sizes, increase latent periods (the time between infection and lysis), or both [1, 3]. These changes in viral traits in response to host state and nutrient condition are examples of phenotypic plasticity, in which behavioral, morphological and physiological traits change in response to changes in the host and/or the environment [4].

The degree of phenotypic plasticity for intracellular viral traits varies across virus-microbe-nutrient systems. Decreases in viral production in response to limitations in essential nutrients like nitrogen and phosphorus are observed across a range of viruses and hosts, from bacteria to phytoplankton [1, 3, 5, 6]. However, viral plasticity is not universal. Phage Paride and phage SEP1 can infect dormant cultures of *Pseudomonas aeruginosa* and *Staphylococcus epidermis* respectively [7, 8, 9]. Viruses that can productively infect and lyse dormant cells typically have large genomes with additional genes that reduce viral dependence on cellular metabolism and intracellular resources [7, 9]. Such decoupling may come with costs. For phage Paride, infection and viral replication in dormant hosts are linked to host stress response genes that may facilitate access to stored resources during cellular starvation – albeit with reduced burst size and longer latent periods. Interestingly, clones of phage Paride passaged on dormant cultures exhibited increased burst size and reduced latent period (i.e higher maturation rate) compared to wildtype Paride, suggesting that plasticity of these viral traits is evolvable [7]. In addition, some viruses appear to be less sensitive than their host cells to certain nutrient limitations: viruses Syn9 and S-PM2 are both able to maintain viral production in cyanobacteria *Synechococcus* when their host’s growth rate decreases due to iron limitation [10]. Viruses may thus respond differently as hosts and environments vary, shaping infection outcomes along with their hosts [11].

Despite evidence for variable maturation rate plasticity across virus-host pairs and environments [1, 3, 5, 6, 7, 8, 9, 10], theoretical studies on plasticity have mostly focused on lysis rate plasticity. Early theory on lysis rate plasticity assumed constant cell density and predicted that the “optimal” latent period decreases with host quality [12, 13]. More recent eco-evolutionary models show similar results in chemostat conditions, finding that the evolutionarily stable latent period decreases and burst size increases with increasing nutrient input and host growth rate, respectively [14]. Alternative models that connect traits with population dynamics have included maturation rate plasticity in terms of a host density-dependent or resource-dependent lysis rate or burst size [15, 16, 17], or by fitting the relationship between viral replication rate and host growth rate to experimental data [14]. However, the extent to which plasticity in intracellular viral traits is adaptively favored given distinct environmental conditions remains unclear.

Here we develop an eco-evolutionary modeling framework to investigate the adaptive benefit of plasticity of intracellular viral traits with a focus on viral maturation and lysis rates. Maturation rate plasticity denotes the degree of dependence of the viral maturation rate on nutrient concentration. Similarly, lysis rate plasticity denotes the dependence of the lysis rate (i.e., inverse of the latent period) on nutrient availability. Together, these maturation rate and lysis rate plasticity parameters dictate changes in burst size in response to nutrient concentration. Throughout, we investigate the potential benefits of plasticity while assuming that decoupling viral traits from host metabolic state comes with a cost (assumed to decrease the maximum attainable maturation rate in nutrient replete conditions). We first use this framework to quantify plasticity from experimental data and find evidence for high maturation rate plasticity and intermediate lysis rate plasticity within studies of viral traits in changing host conditions. We then use a combination of theory and eco-evolutionary simulations to identify a broad range of conditions where maturation and lysis rate plasticity maximize invasion fitness and viral growth. Finally we explore how changes in nutrient concentrations and host and viral abundances can facilitate the coexistence of phages with different degrees of plasticity, even when competing for a shared host population.

## Methods

### Viral trait plasticity

The maturation rate, *m*(*R*), and lysis rate, *η*(*R*), are modeled as linear functions of the host’s relative growth rate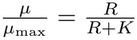, which itself is a saturating Monod function of the resource concentration *R*. The parameters *ρ*_*m*_ and *ρ*_*η*_ modulate maturation rate and lysis rate plasticity, respectively (Figure 1A,B). The burst size *β*(*R*) follows from *m*(*R*) and *η*(*R*) (Figure 1C,E):

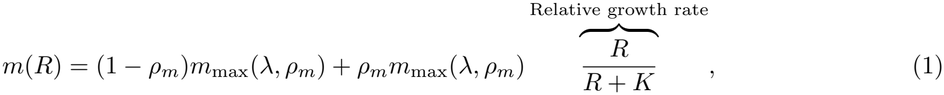

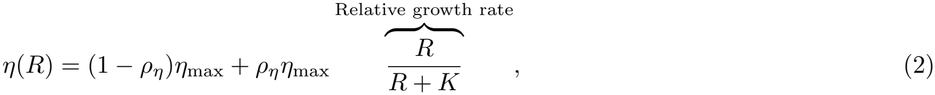

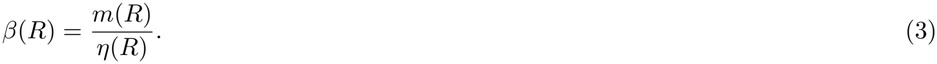

**Figure 1:**
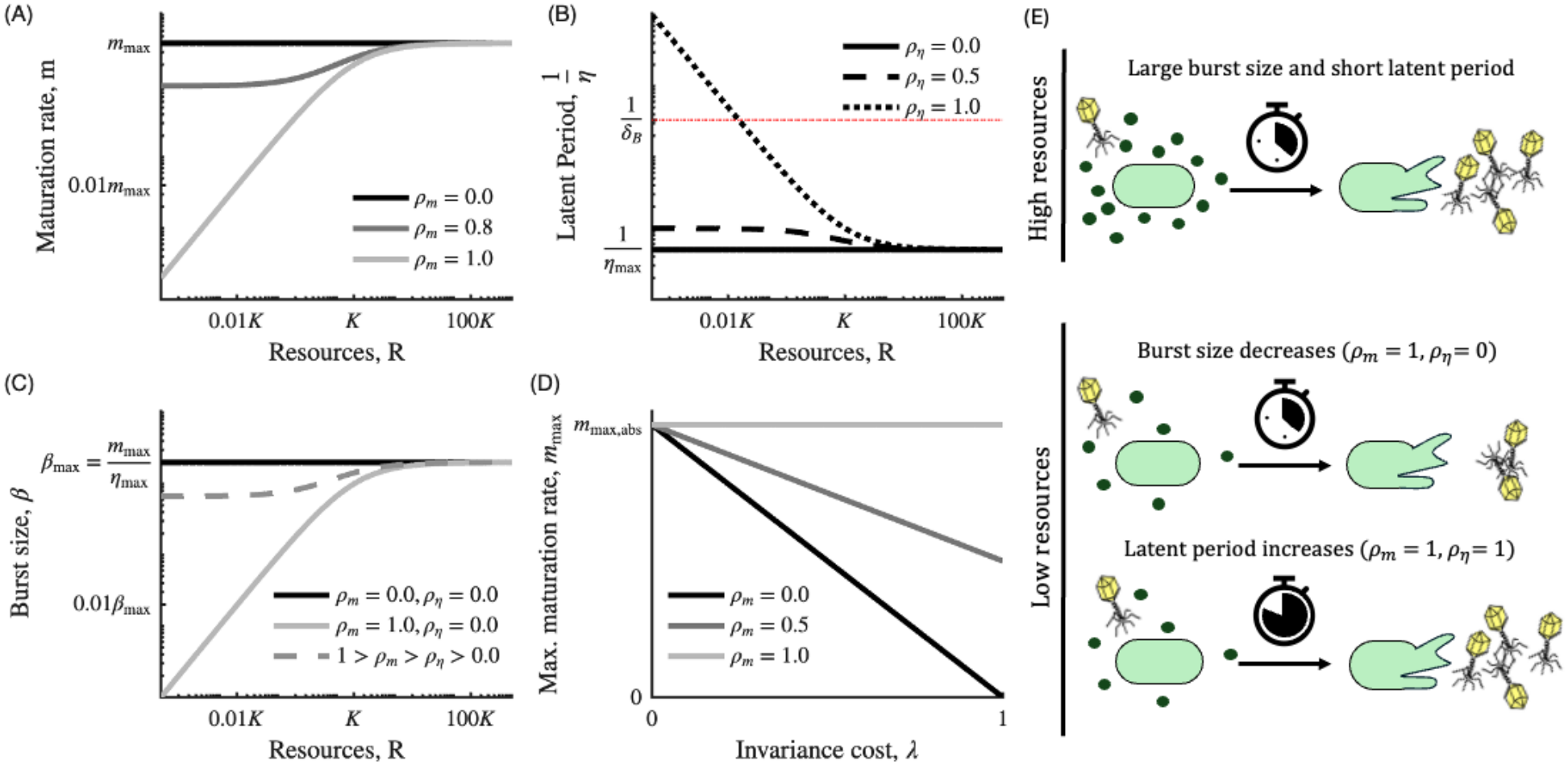
Model description. **(A)** The maturation rate *m* increases with resources, from a minimum determined by maturation plasticity *ρ*_*m*_ to its maximum *m*_max_. **(B)** The latent period 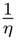 decreases from a maximum determined by lysis rate plasticity *ρ*_*η*_ to its minimum 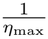 . The mean life span of an infected Cell 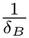 is shown with a dotted red line. **(C)** The burst size is determined by the maturation rate and latent period. Identical maturation rate and lysis rate plasticity values results in a constant burst size at its maximum 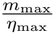. When lysis rate plasticity *ρ*_*η*_ is lower than maturation rate plasticity *ρ*_*m*_, burst size decreases as R decreases. **(D)** The maximum maturation rate decreases as invariance cost increases, with maturation rate plasticity *ρ*_*m*_ determining the slope. Invariant viruses have a null maximum maturation rate if the invariance cost is 1, while fully plastic viruses maintain a maximum maturation rate *m*_max,abs_ for all invariance costs. **(E)** Visual summary of the effect of maturation rate plasticity *ρ*_*m*_ and lysis rate plasticity *ρ*_*η*_ on burst size and latent period in low- and high-resource environments.

For trait invariant viruses (*ρ*_*m*_ = *ρ*_*η*_ = 0), the maturation rate *m*(*R*) and lysis rate *η*(*R*) are not affected by *R* and are always equal to their maxima *m*_max_(*λ, ρ*_*m*_) and *η*_max_. On the other hand, the maturation rate of maximally plastic viruses (*ρ*_*m*_ = 1) is proportional to the relative growth rate of the host and decreases to 0 when *R* is 0. For all other values of *ρ*_*m*_ and *ρ*_*η*_, the maturation rate *m*(*R*) and lysis rate *η*(*R*) reach their maxima *m*_max_(*λ, ρ*_*m*_) and *η*_max_, respectively, when *R* ≫ *K*. When *R* = 0, the maturation rate *m* and lysis rate *η* take a positive value if 0 ≤ *ρ*_*m*_, *ρ*_*η*_ *<* 1, representing a non-zero maturation and lysis rate in non-growing bacteria. Plasticity estimates above 1 lead to a more-than-proportional decrease in traits as resources decrease, resulting in the maturation rate or lysis rate decreasing to 0 for a non-zero positive resource concentration *R*.

We assume that the lysis rate can only change to accommodate at most a proportional change in the maturation rate in response to resource concentration; therefore, we limit our analysis to the parameter space *ρ*_*η*_ *< ρ*_*m*_ (Figure S1). This means that viruses can only lengthen their latent period in response to a decrease in maturation rate. At the extreme, a virus with a fully invariant maturation rate (*ρ*_*m*_ = 0) must have a fully invariant lysis rate (*ρ*_*η*_ = 0) - modifications of these assumptions are revisited in the Discussion.

A tradeoff between maintaining a maturation rate independent of host physiology and the maximum possible maturation rate (i.e., what we term ‘trait invariance’) is modeled by linking the maximum maturation rate *m*_max_(*λ, ρ*_*m*_) to the trait invariance cost *λ* (Figure 1D):

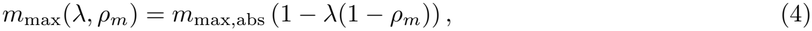

where *m*_max,abs_ is the absolute maximum maturation rate. An invariance cost *λ* of 0 corresponds to no replication cost for invariant viruses, while *λ* = 1 results in a maximum maturation rate *m*_max_ of 0 for invariant viruses (i.e., invariant viruses cannot replicate if cost *λ* = 1). We do not assume a maturation rate benefit for viruses with *ρ*_*m*_ *>* 1. We thus restrict the values that *ρ*_*m*_ and *ρ*_*η*_ can take in simulations between 0 and 1, spanning the range of the tradeoff described here.

#### Estimation of plasticity parameters from experimental data

Experimental studies on viral plasticity were obtained from Google Scholar using any combination of the keywords ‘growth’, ‘virus’, ‘infection’, ‘nutrient limitation’, and ‘plasticity’. Data were then included in this analysis if host growth rate, latent period and burst size had been measured for at least two different host growth conditions. We always calculated the maturation rate from the latent period and burst size (*m* = *βη*), as the eclipse period was not always provided and was not included in our model.

Many studies measured viral traits at two growth rates only, typically in a nutrient-limited and a nutrient-replete state. Plasticity estimates *ρ*_*m*_ and *ρ*_*η*_ were calculated from Eqs. 1 and 2, with 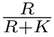 expressed as the relative growth rate 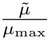, where 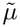 is the observed growth rate. We first assume that *µ*_max_ and *m*_max_ can be estimated directly as the growth rate and maturation rate in nutrient-replete conditions. We then estimate *ρ*_*m*_ from the observed maturation rate 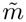 and observed host growth rate 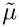 in nutrient-limited conditions:

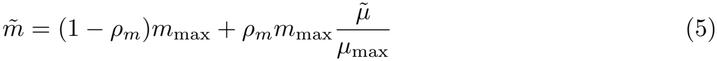

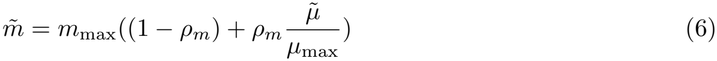

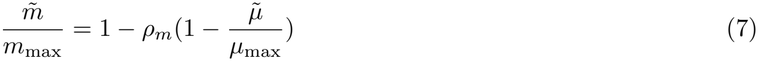

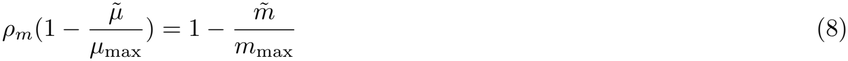

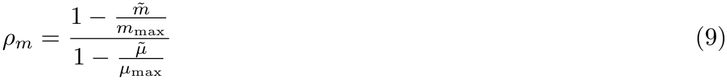

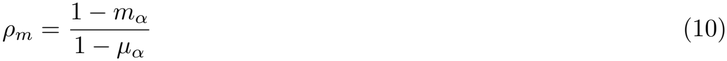

where *m*_*α*_ and *µ*_*α*_ are the relative maturation rate and bacterial growth rate, respectively, in nutrient-limited compared to nutrient-replete conditions. We similarly derive the lysis rate *ρ*_*η*_:

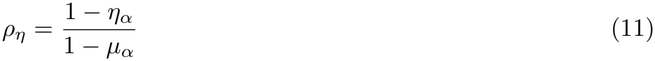

where *η*_*α*_ is the relative lysis rate in nutrient-limited compared to nutrient-replete conditions.

Three studies of model phages measured viral traits across a range of growth rates, allowing for statistical fitting of *ρ*_*m*_ and *ρ*_*η*_. We used the optim function in R [18] to find the least square estimate of *ρ*_*m*_ and *ρ*_*η*_, assuming *µ*_max_ to be the highest growth rate observed in the study and *m*_max_ to be the maturation rate at that growth rate. Note that we did not restrict the possible values of *ρ*_*m*_ and *ρ*_*η*_ in these analyses.

#### RSInV Model

We develop a two-stage chemostat model of virus-host dynamics, with inflow of resources *R* and susceptible cells *S*, and outflow of resources, cells (including infected cells *I*) and viruses *V*, following [19]. The growth rate of susceptible cells is a Monod function of resources. Adsorption of viruses to hosts is assumed to follow mass-action kinetics. We include *n* infected stages to account for variability in latent period [20, 21, 22, 23]. This linear-chain trick method results in an Erlang distributed latent period with mean 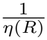 and coefficient of variation 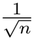 [21]. Infected cells lyse after progressing through the infected stages, releasing progeny that can infect other susceptible cells, decay, or wash out:

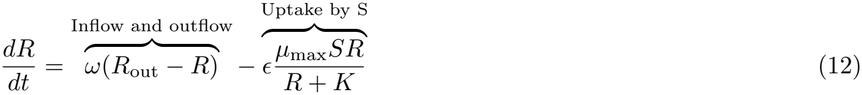

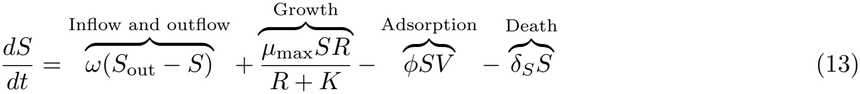

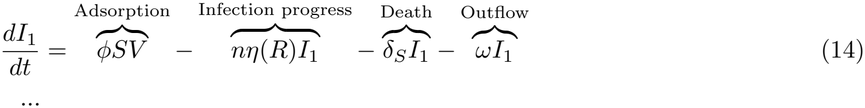

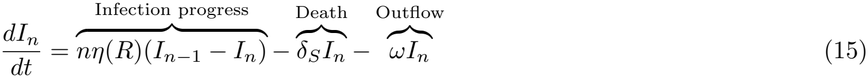

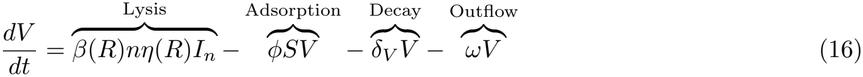

where *ω* is the mixing rate, *R*_out_ and *S*_out_ are the outside concentrations of resources and susceptible cells respectively, *ϵ* is the conversion rate between resources and cells, *µ*_max_ is the maximum cell growth rate, *ϕ* is the adsorption rate, *δ*_*S*_ is the death rate for all cells, *δ*_*V*_ is the viral decay rate, *η*(*R*) is the resource-dependent lysis rate and *β*(*R*) is the resource-dependent burst size. We assume that virions do not adsorb to infected cells.

#### Invasion criteria

We evaluate the effect of maturation rate plasticity *ρ*_*m*_ and lysis rate plasticity *ρ*_*η*_ on the basic reproduction number, *ℛ*_0_, of viruses – the number of progeny viruses produced on average by a single infectious virion in its lifetime in an otherwise susceptible host population [24]. *ℛ*_0_ is equal to the product of the probability that the virus survives until adsorption 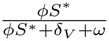, the probability that the infected cell survives through the *n* infected stages until lysis 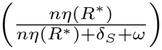and the burst size *β*(*R*^*\**^), where *R*^*\**^ and *S*^*\**^ are the resource and susceptible cell density at equilibrium:

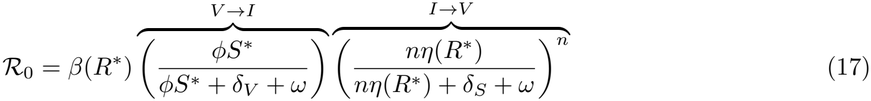

where *η*(*R*^*\**^) and *β*(*R*^*\**^) depend on the equilibrium resource concentration *R*^*\**^, modulated by *ρ*_*m*_ and *ρ*_*η*_, as described above (Eq. 1-3).

#### Viral growth rate at invasion

We calculate an effective viral growth rate assuming constant host density [12]. Under this assumption, if each virion produced *ℛ*_0_ offspring per infection cycle and the average duration of each infection cycle is *L*_*r*_, then the virion population at time *t* is given by

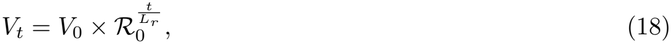

where *V*_*t*_ is the virion population at time *t* and *V*_0_ is the initial virion population. 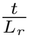 is simply the number of infection cycles over time *t*.

The average duration of an infection cycle is the sum of the average latent period and the average time it would take for a new virion offspring to find a susceptible host:

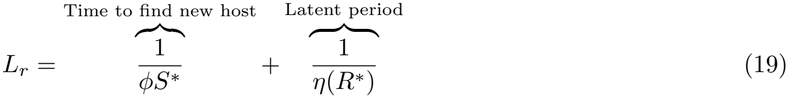

Comparing Equation 18 to the standard exponential growth equation (*x*_*t*_ = *x*_0_*e*^*rt*^), we obtain the viral growth rate:

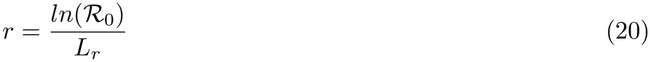

#### Competition simulations

A number *m* of competing viruses are introduced to a system with resources and susceptible cells at their virus-free equilibrium. All viruses are introduced at the same time at initial densities of 10^7^ *L*^*−*1^ per virus type. We assume that cells infected by one virus cannot be infected by another (i.e. superinfection exclusion) and that viruses only differ in their plasticity, resulting in differences in their lysis rate *η*_*m*_(*R*) and burst size *β*_*m*_(*R*):

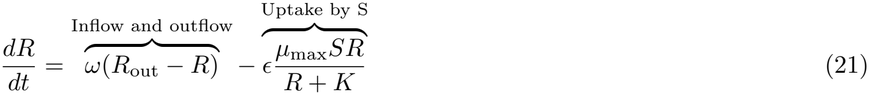

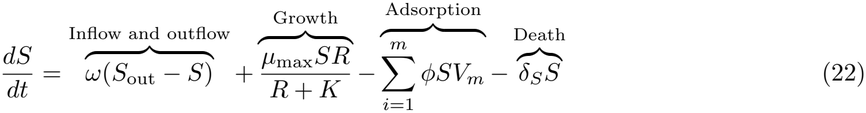

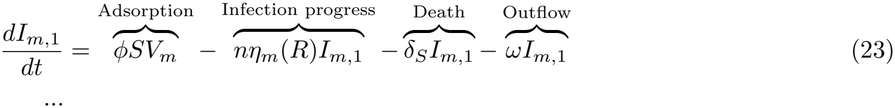

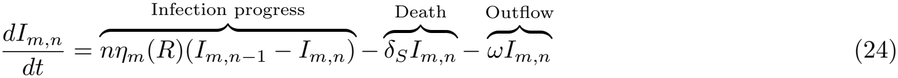

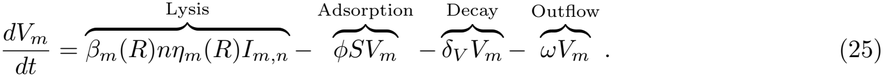

Here, *I*_*m,n*_ denotes the density of cells infected by virus-type *m* that are currently in infected state *n*.

#### Numerical methods

We simulate model dynamics over time using the solver ode45 in Matlab2025b [25]. In competition simulations with constant resource inflow, viral plasticity traits resulting in the highest viral density at the end of the simulation are considered optimal. In competition simulations with varying mixing rate, a virus or host is considered extinct when its density drops below 10^*−*7^ L^*−*1^. We define coexistence as the ability of virus to mutually invade each other from a density of 1 L^*−*1^ when the other virus as at steady state.

#### Sensitivity analysis

We performed a sensitivity analysis using a Latin Hypercube Sampling design for the virus-free equilibrium for mixing rate *ω*, outside resource density *R*_out_, outside cell density *S*_out_, maximum cell growth rate *µ*_max_ and half-saturation constant *K*. Virus-free equilibrium values were determined analytically. When viruses were included, we varied parameters one-by-one and obtained equilibrium values through 1000-day numerical simulations for mixing rate *ω*, outside resource density *R*_out_, maximum cell growth rate *µ*_max_, adsorption rate *ϕ*, absolute maximum maturation rate *m*_max,abs_, and maximum lysis rate *η*_max_. We parametrized the model for phytoplankton and their viruses. Default values are shown in Table 1.

**Table 1:** Parameter descriptions and values for phytoplankton and their viruses.

| **Description** | **Parameter** | **Value** | **Unit** | **References** |
|---|---|---|---|---|
| Mixing rate | *ω* | 0.1 | d<sup>−1</sup> | [16] |
| External resource density | *R*<sub>out</sub> | 100 | *n*mol L<sup>−1</sup> |  |
| External cell density | *S*<sub>out</sub> | 10<sup>7</sup> | cells L<sup>−1</sup> | [19] |
| Cell nutrient quota | *ϵ* | 6.4 × 10<sup>−8</sup> | *n*mol cell<sup>−1</sup> | [16] |
| Max. cell growth rate | *μ*<sub>*max*</sub> | 1 | d<sup>−1</sup> | [16] |
| Half-saturation constant | K | 20 | *n* mol L<sup>−1</sup> | [16] |
| Adsorption rate | *ϕ* | 7 × 10<sup>−9</sup> | L cell<sup>−1</sup> d<sup>−1</sup> | [19, 26] |
| Max. replication rate | *m*<sub>*max,abs*</sub> | 400 | d<sup>−1</sup> | [16] |
| Max. lysis rate | *η*<sub>*max*</sub> | 2 | d<sup>−1</sup> | [16] |
| No. of infected classes | *n* | 20 |  | [21] |
| Invariance cost | *λ* | 0-1 |  |  |
| Replication rate plasticity | *ρ*<sub>*m*</sub> | 0-1 |  |  |
| Lysis rate plasticity | *ρ*<sub>*η*</sub> | 0-1 |  |  |
| Cell death rate | *δ*<sub>*S*</sub> | 0.03-0.3 | d<sup>−1</sup> | [16] |
| Viral decay rate | *δ*<sub>*V*</sub> | 0.5 | d<sup>−1</sup> | [16] |

## Results

### Empirical estimates of plasticity in maturation rate and lysis rate

We first compiled data from the literature and estimated the maturation rate plasticity (*ρ*_*m*_) and lysis rate plasticity (*ρ*_*η*_) (Table 2, Figure 2). With the exception of viruses in iron-limited conditions, we find that viruses subject to variable nutrient conditions exhibit high maturation rate plasticity and lower lysis rate plasticity, ranging from 0.25 to 0.75 (Table 2). This combination of high maturation rate plasticity and intermediate lysis rate plasticity leads to moderate plasticity in predicted burst size (Figure S2). Notably, maturation rate plasticity estimates include some values above 1, indicating a decrease in maturation rate larger than a proportional decrease to the host’s growth rate.

**Table 2:**
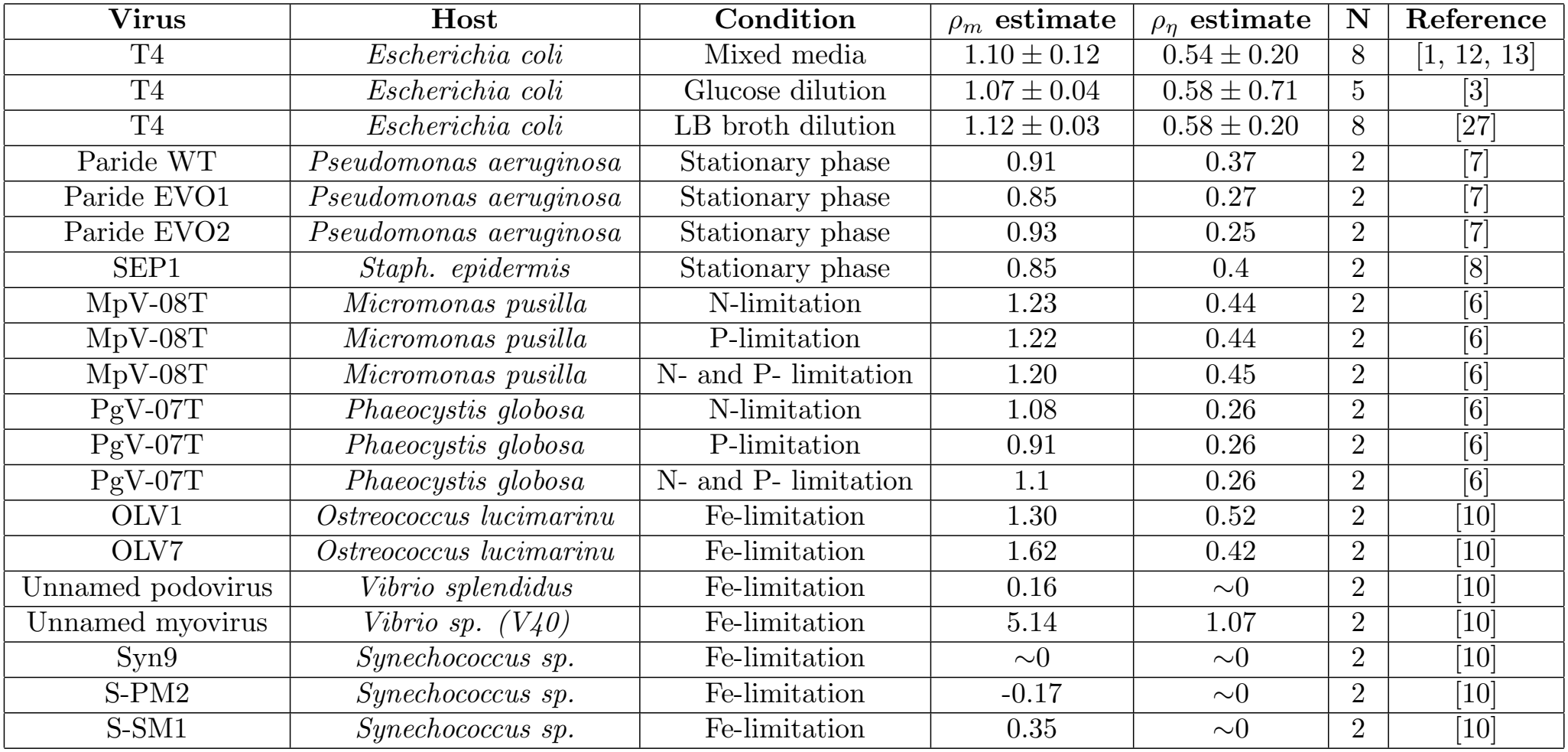
Estimates for *ρ*_*m*_ and *ρ*_*η*_ in experimental data. Number of samples per experiment is given by N.

| **Virus** | **Host** | **Condition** | ρ<sub>*m*</sub> **estimate** | ρ<sub>*η*</sub> **estimate** | **N** | **Reference** |
|---|---|---|---|---|---|---|
| T4 | *Escherichia coli* | Mixed media | 1.10 ± 0.12 | 0.54 ± 0.20 | 8 | [1, 12, 13] |
| T4 | *Escherichia coli* | Glucose dilution | 1.07 ± 0.04 | 0.58 ± 0.71 | 5 | [3] |
| T4 | *Escherichia coli* | LB broth dilution | 1.12 ± 0.03 | 0.58 ± 0.20 | 8 | [27] |
| Paride WT | *Pseudomonas aeruginosa* | Stationary phase | 0.91 | 0.37 | 2 | [7] |
| Paride EVO1 | *Pseudomonas aeruginosa* | Stationary phase | 0.85 | 0.27 | 2 | [7] |
| Paride EVO2 | *Pseudomonas aeruginosa* | Stationary phase | 0.93 | 0.25 | 2 | [7] |
| SEP1 | *Staph. epidermis* | Stationary phase | 0.85 | 0.4 | 2 | [8] |
| MpV-08T | *Micromonas pusilla* | N-limitation | 1.23 | 0.44 | 2 | [6] |
| MpV-08T | *Micromonas pusilla* | P-limitation | 1.22 | 0.44 | 2 | [6] |
| MpV-08T | *Micromonas pusilla* | N- and P- limitation | 1.20 | 0.45 | 2 | [6] |
| PgV-07T | *Phaeocystis globosa* | N-limitation | 1.08 | 0.26 | 2 | [6] |
| PgV-07T | *Phaeocystis globosa* | P-limitation | 0.91 | 0.26 | 2 | [6] |
| PgV-07T | *Phaeocystis globosa* | N- and P- limitation | 1.1 | 0.26 | 2 | [6] |
| OLV1 | *Ostreococcus lucimarinu* | Fe-limitation | 1.30 | 0.52 | 2 | [10] |
| OLV7 | *Ostreococcus lucimarinu* | Fe-limitation | 1.62 | 0.42 | 2 | [10] |
| Unnamed podovirus | *Vibrio splendidus* | Fe-limitation | 0.16 | ∼0 | 2 | [10] |
| Unnamed myovirus | *Vibrio sp. (V40)* | Fe-limitation | 5.14 | 1.07 | 2 | [10] |
| Syn9 | *Synechococcus sp.* | Fe-limitation | ∼0 | ∼0 | 2 | [10] |
| S-PM2 | *Synechococcus sp.* | Fe-limitation | -0.17 | ∼0 | 2 | [10] |
| S-SM1 | *Synechococcus sp.* | Fe-limitation | 0.35 | ∼0 | 2 | [10] |

**Figure 2:**
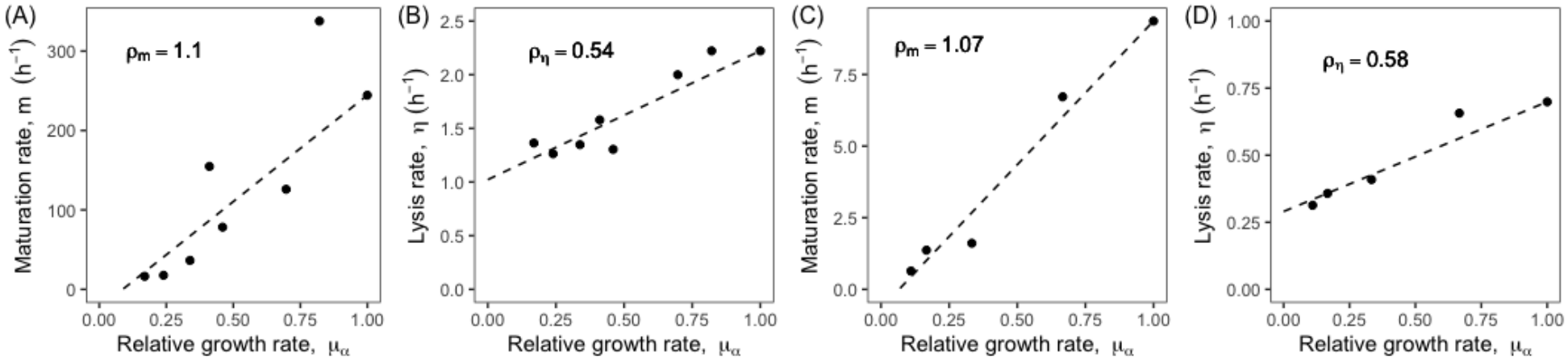
Maturation rate plasticity (*ρ*_*m*_) and lysis rate plasticity (*ρ*_*η*_) estimated from published experimental data for phage T4. Parameters *ρ*_*m*_ and *ρ*_*η*_ were estimated by fitting Equations 1 and 2 to latent period and burst size measurements at different bacterial growth rates. Maturation rates (A, C) and lysis rates (B, D) for phage T4 and *E. coli* grown in different media including LB broth and supplemented minimal media (A, B) [1], and dilution rates (corresponding to growth rates) of 0.3, 0.2, 0.1, and 0.033 *h*^*−*1^ (C, D) [3]. Dotted lines show best-fit relationships for *ρ*_*m*_ and *ρ*_*η*_ estimated through the least square error method.

We observed extreme values and greater variability in plasticity across virus-host pairs in iron-limited environments. While some viruses exhibited maturation rate plasticity values above 1, others showed no maturation plasticity (*ρ*_*m*_ close to or equal to 0). We observed similar variability in lysis rate plasticity as well. Finally, both maturation rate and lysis rate plasticity values were found to decrease when phage Paride was passaged on dormant bacterial populations([7], Table 2), suggesting adaptation to poor host conditions. Despite these variations across nutrient conditions and virus-host pairs, lysis rate plasticity estimates were always lower than maturation rate plasticity estimates. These findings provide expanded quantitative evidence that maturation rate and lysis rates are plastic, and that the degree of plasticity varies with virus-host systems subject to distinct limiting nutrients.

### High lysis rate plasticity enhances viral invasion when resources are abundant

We consider how viral trait plasticity affects viral invasion of a susceptible bacterial population by calculating the basic reproduction number *ℛ*_0_ and varying maturation rate and lysis rate plasticity separately. When resources are abundant, we find that viruses with invariant traits, *ρ*_*m*_ = 0, maximize *ℛ*_0_, in the absence of invariance costs (Figure 3A). However, with a positive invariance cost, higher values of *ρ*_*m*_ maximize *ℛ*_0_ given sufficiently high resource levels (Figure 3B). Assuming a fully plastic maturation rate (*ρ*_*m*_ = 1), we find that the lysis rate plasticity that maximize *ℛ*_0_ also increases with resource concentration 3C).

**Figure 3:**
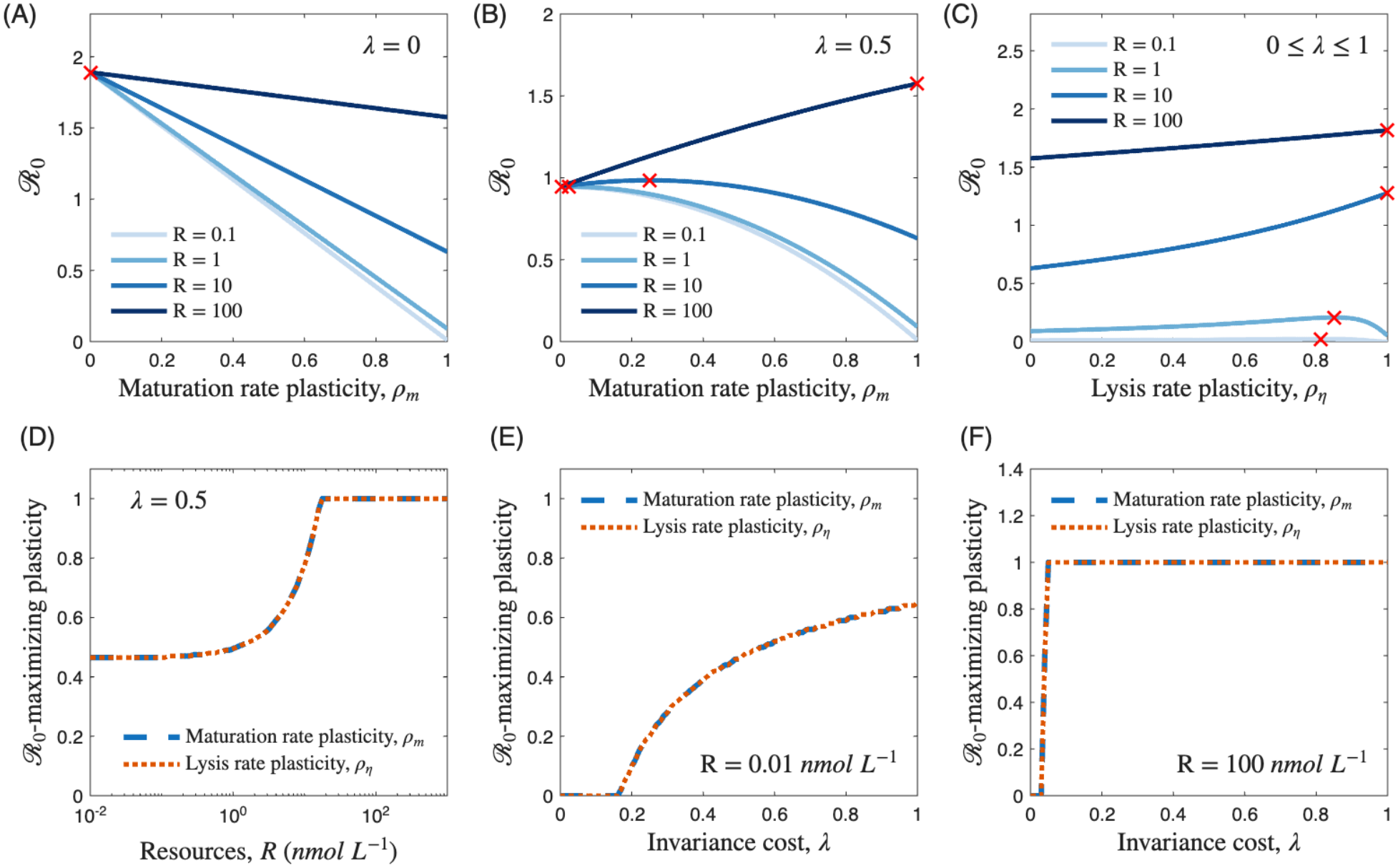
Viral plasticity maximizing *ℛ*_0_ at fixed susceptible cell and resource density. Viral reproduction number *ℛ*_0_ across maturation rate plasticity *ρ*_*m*_ at four resource concentrations R (*nmol L*^*−*1^) for invariance cost *λ* = 0 **(A)** and *λ* = 0.5 **(B)**. Lysis rate plasticity was set to 0. The maximum *ℛ*_0_ for each resource concentration is indicated with a red cross. Susceptible cell density was set to *S* = 10^6^*L*^*−*1^. **(C)** Viral reproduction number *ℛ*_0_ across lysis rate plasticity *ρ*_*η*_, keeping maturation rate plasticity constant at 1. The pair of maturation rate and lysis rate plasticity that maximize *ℛ*_0_ was then calculated across resource concentration R (*λ* = 0.5) **(D)** and across invariance cost (*R* = 0.01 *nmol L*^*−*1^ or *R* = 100 *nmol L*^*−*1^) **(E**,**F)**.

We then calculate combinations of maturation rate plasticity *ρ*_*m*_ and lysis rate plasticity *ρ*_*η*_ that maximize R_0_ across changing levels of susceptible host densities, resource densities, and invariance costs. We find that the plasticity *ρ*_*m*_ and *ρ*_*η*_ that maximize *ℛ*_0_ are independent of the density of susceptible cells and equal to each other (Figure S3). In other words, any decrease in maturation rate results in an increase in latent period rather than a decrease in burst size. Recall that *ℛ*_0_ depends linearly on *β* and saturates with increasing *η* (Eq. 17). Reducing the lysis rate *η* has a much smaller effect on *ℛ*_0_ than a proportional decrease in burst size *β*, as long as lysis rates are much faster than washout and decay rates (which is the case in most systems). The maturation rate plasticity *ρ*_*m*_ that maximizes *ℛ*_0_ increases with resource concentration (Figure 3D), as plasticity becomes less limiting when resource concentration increases.

Invariance costs increase the plasticity *ρ*_*m*_ and *ρ*_*η*_ that maximize *ℛ*_0_ at low resource densities (Figure 3E). Similarly at high resource densities, high maturation rate and lysis rate plasticity maximize R_0_ for a wide range of invariance costs. Low maturation rate plasticity only maximizes *ℛ*_0_ for very low values of invariance costs that are unlikely to apply to virus-host-nutrient systems (Figure 3F).

### Low lysis rate plasticity enhances viral invasion when mixing rate is high

We next ask how *ℛ*_0_-maximizing plasticity varies across resource and susceptible cell densities driven by underlying parameter changes. When holding parameters constant, we observed that the plasticity *ρ*_*m*_ and *ρ*_*η*_ that maximize *ℛ*_0_ increase with resource density and do not change with susceptible cell density (Figure 3D, Figure S3). These patterns do not hold when resources and susceptible cell densities vary in response to mixing rate (Figure 4) or other parameters (Figure S4, Figure S5). In a two-stage chemostat, the mixing rate drives both resource and cell densities. Resource concentration increases with mixing rate, before reaching a plateau (Figure 4A, Figure S5). Susceptible cell density first increases before decreasing sharply as mixing rate increases (Figure 4B).

**Figure 4:**
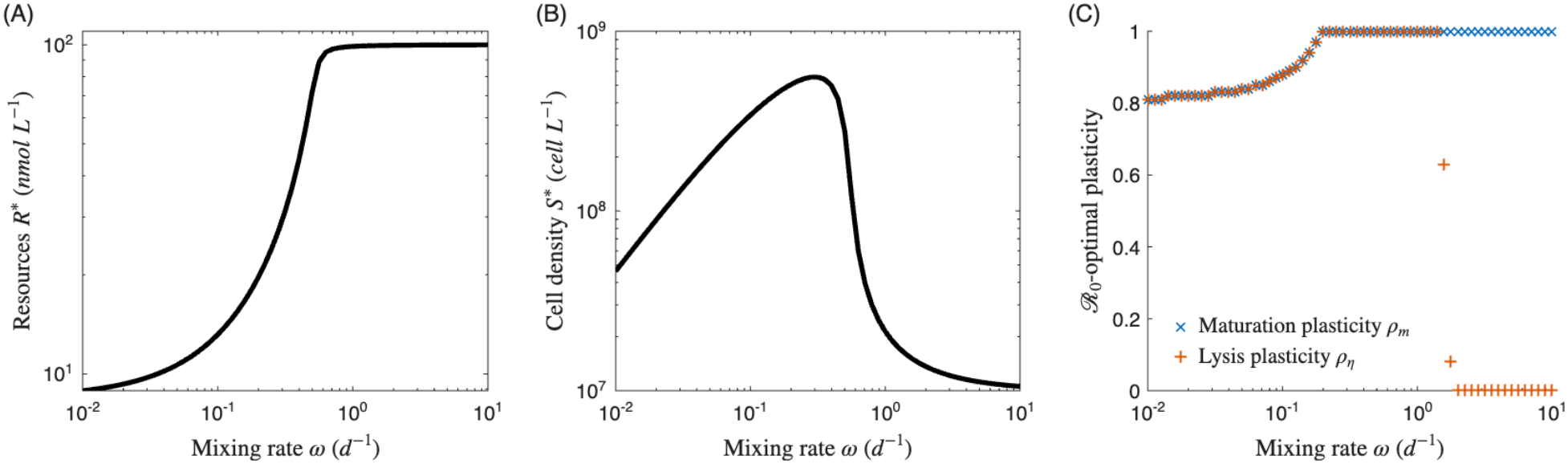
*ℛ*_0_-maximizing plasticity across mixing rate *ω*. **(A)** Equilibrium resource concentration *R*^*\**^ across mixing rate *ω*. **(B)** Equilibrium susceptible host density *S*^*\**^ across mixing rate *ω*. **(C)** Maturation rate plasticity *ρ*_*m*_ and lysis rate plasticity *ρ*_*η*_ that maximize *ℛ*_0_ at equilibrium across mixing rate *ω*. Invariance cost *λ* is set to 0.5.

The combination of increasing resources and susceptible cell density first favors increasing maturation rate and lysis rate plasticity as mixing rate increases (Figure 4C). As the mixing rate continues to increase, *ρ*_*m*_ and *ρ*_*η*_ that maximize *ℛ*_0_ are both maintained at 1. Finally, very high mixing rates lead to a sharp decrease in the density of susceptible cells. This decline drives the maturation rate plasticity *ρ*_*η*_ that maximizes *ℛ*_0_ to 0, effectively creating plasticity in the burst size. High mixing rates result in the removal of infected cells from the system before lysis, which favors viruses with low lysis rate plasticity.

Besides mixing rate, we find that the inflowing resource concentration and the maximum cell growth rate also affect invasion-optimal plasticity. Inflowing resource concentration has a more direct effect on *ℛ*_0_-maximizing *ρ*_*m*_ and *ρ*_*η*_; both increase with inflowing resource concentration as resource density in the chemostat increases in response (Figure S4, Figure S5). Finally, we find that increases in the maximum cell growth rate past a certain threshold (∼ 1 *d*^*−*1^) leads to a decrease in *ρ*_*m*_ and *ρ*_*η*_ that maximize *ℛ*_0_, as this increase in growth rate leads to a decrease in resource concentration (Figure S4, Figure S5).

### Burst size plasticity maximizes viral growth rate when hosts are replete

We next asked how viral plasticity affects the growth rate of the viral population during and shortly after invasion of a naive, susceptible cell population. Unlike for *ℛ*_0_-maximization (Figure S3), growth-maximizing plasticity *ρ*_*m*_ and *ρ*_*η*_ vary with the density of susceptible cells *S* (Figure 5A,B). When host density is low, the optimal lysis rate *ρ*_*η*_ matches the optimal maturation rate plasticity *ρ*_*m*_, and both decrease with host density and increase with resources. As host density increases, we observe a shift at high cell density and resources: viruses with the highest growth have a plastic maturation rate but an invariant lysis rate (equivalent to a plastic burst size). In other words, viruses that lyse early grow faster when host density is high. Finally, we ask how invariance cost affects growth-maximizing plasticity. At low host density, fully plastic viruses have the highest growth for any invariance cost above 0.4 (Figure 5C). On the other hand, high host density results in a lower maturation rate plasticity and a lower lysis rate plasticity that maximize growth for a given cost (Figure 5D). Overall, viral population invasion growth rates are maximized by high lysis rate plasticity at low densities of susceptible cells and by high burst size plasticity at high densities of susceptible cells; as before, increases in the cost of invariance tend to increase the adaptive value of maturation rate plasticity.

**Figure 5:**
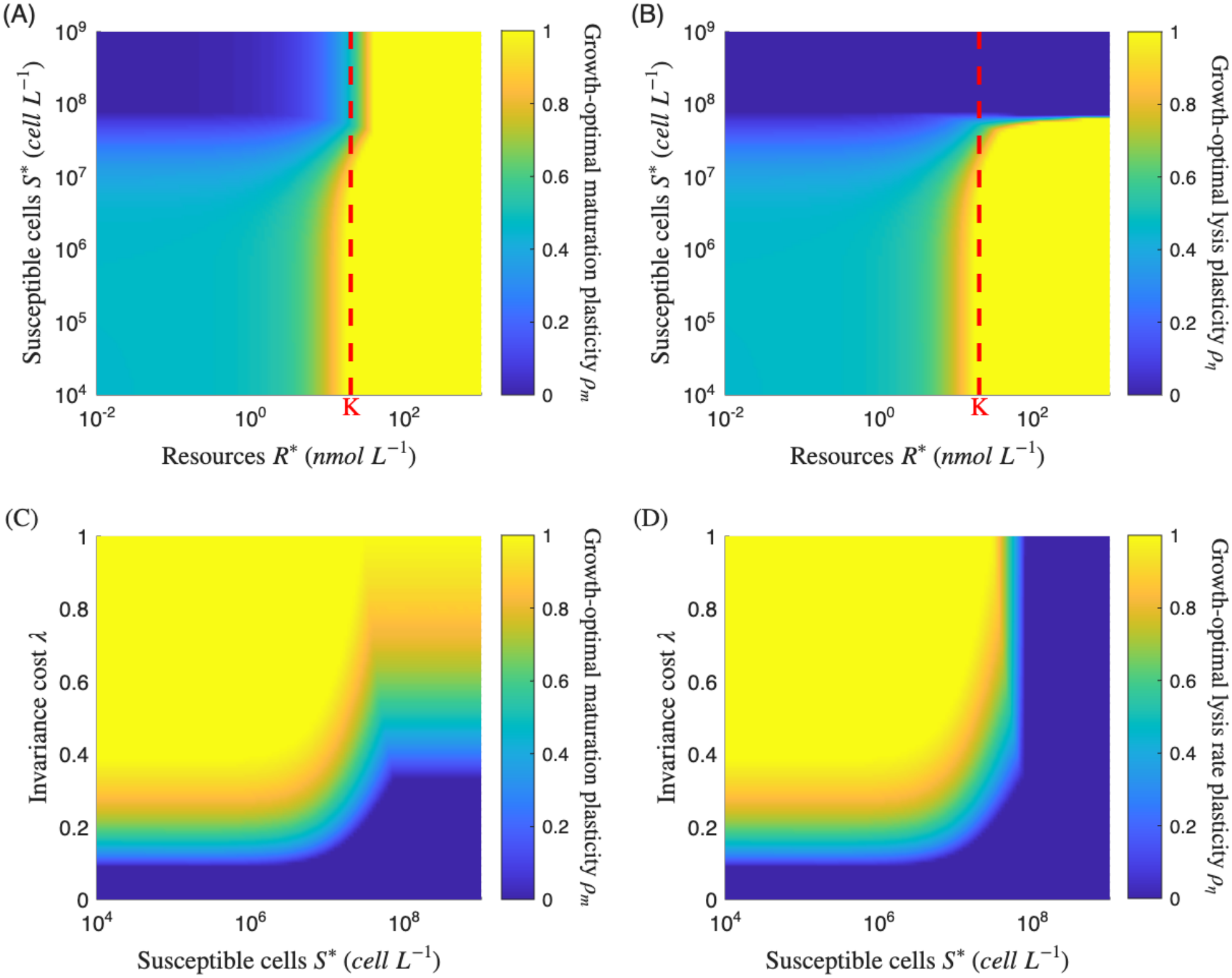
Growth-maximizing plasticity for viral growth at invasion. The pair of maturation rate and lysis rate plasticity that maximizes viral growth shortly after invasion (assuming a fixed host pool) was calculated across resources and susceptible cells **(A**,**B)** and across susceptible cells and invariance cost **(C**,**D)**. Invariance cost *λ* is set 0.5 and resources are set to *K* in the top and bottom panels respectively. The red dashed line shows *R* = *K*.

### Optimal plasticity in viral competition

Thus far, we have shown how plasticity in viral traits impacts individual and population level metrics of viral invasion in fixed conditions. However, evolutionary outcomes are shaped by competition in environments with non-linear feedbacks between host, viral and resource density. In order to assess the relationship between invasion and long-term outcomes, we simulate competitions between viruses with different maturation rate and lysis rate plasticity infecting a single population of susceptible hosts. Optimal *ρ*_*m*_ and *ρ*_*η*_ were determined as the maturation rate plasticity and lysis rate plasticity of the virus with the highest density long after a transient period, here set to 3000 days. We varied parameters that have the largest influence on resource and susceptibility density (Figure S6): inflowing resource density *R*_out_, adsorption rate *ϕ*, mixing rate *ω*, maximum lysis rate *η*_*max*_, and the maximum maturation rate *m*_max,abs_.

In competition simulations, optimal maturation rate plasticity *ρ*_*m*_ and lysis rate plasticity *ρ*_*η*_ were high across the range of adsorption rates and growth rates (Figure S7). As for *ℛ*_0_, we found that optimal *ρ*_*η*_ decreases for very high mixing rate, as bacterial cells are removed quickly from the environment, favoring viruses with shorter latent periods (Figure S7). Low resource density from low inflowing resource concentrations also lowers the optimal *ρ*_*m*_ and *ρ*_*η*_ (Figure S7). Finally, we observed a decrease in optimal *ρ*_*m*_ and *ρ*_*η*_ for lower maximum lysis rate values (Figure S7). A decrease in maximum lysis rate is associated with a decrease in susceptible cell density (Figure S6), which we expect to favor viruses with lysis rate plasticity (Figure 5B). We observe the opposite (Figure S7), suggesting that burst size plasticity (low lysis rate plasticity) is optimal due to direct effects of the maximum lysis rate on viral competition rather than indirect effect through decreased susceptible cell densities.

### Intermittent poor conditions favor viruses with high plasticity

We hypothesize that maturation rate and lysis rate plasticity increase fitness in variable environments by slowing down lysis and viral production when nutrient conditions are poor and susceptible cells are relatively rare. We simulated environments with intermittent periods of low mixing rates, leading to drops in resource and host density. We set the invariance cost to zero, which results in plastic viruses having a lower growth rate than invariant viruses during high-resource periods (Figure 6A). During periods with near-zero resource density, infected cells of viruses with a highly plastic latent period do not lyse and survive for longer while invariant viruses continue lysing infected cells, releasing virus particles that decay relatively quickly (Figure 6A). This difference in lysis rates can result in the survival of plastic viruses but not invariant viruses, given low-nutrient periods that end after extinction of the invariant infected cells but before the extinction of plastic infected cells (Figure 6A).

**Figure 6:**
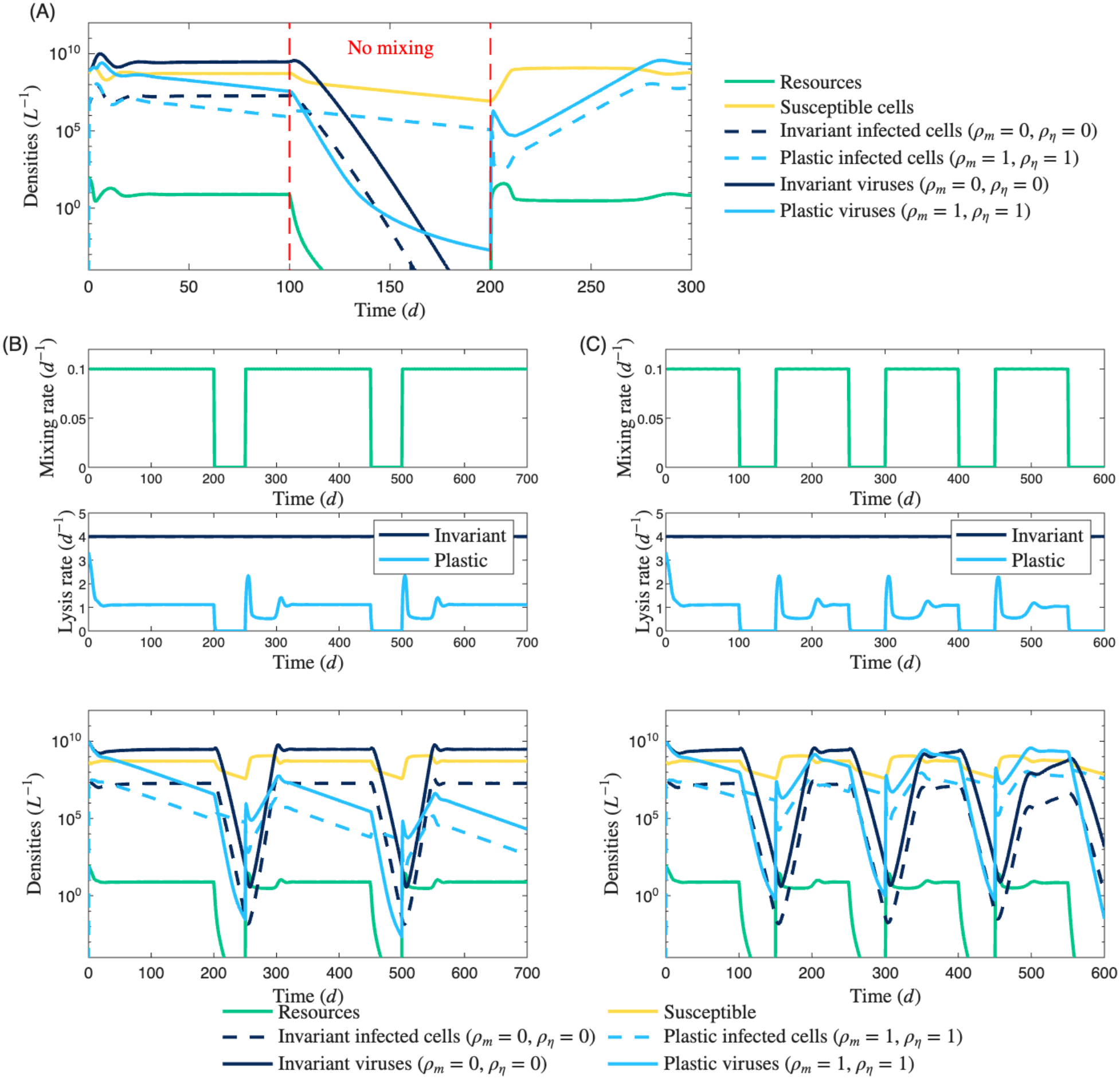
Optimal plasticity in competition simulations in a variable environment. Invariance cost *λ* was set to 0 for these simulations. **(A)** Example dynamics with a drop in mixing from day 100 to 200, causing a drop in resources and lysis of cells infected by a plastic virus over the same period. Invariant viruses go extinct while plastic viruses maintain a high density of infected cells that produce viruses when mixing and resources increase on day 200. **(B)** Plastic viruses decrease in abundance when low-resource periods are infrequent. Mixing rate, lysis rates and densities are shown for a simulation with high-resource and low-resource periods lasting 200 and 50 days respectively. **(C)** Invariant viruses decrease in abundance when low-resource periods are frequent. Mixing rate, lysis rates and densities are shown for a simulation with high-resource and low-resource periods lasting 100 and 50 days respectively. Non-default parameter values are as follows: *ϕ* = 5 × 10^*−*11^ L cell^*−*1^ d^*−*1^, *η*_*max*_ = 4 *d*^*−*1^, *m*_max,abs_ = 100 *d*^*−*1^, *δ*_*S*_ = 0.03 *d*^*−*1^.

We simulated a low-mixing period duration of 50 days, allowing survival of both viruses if starting from the same high density. Infrequent instances of resource-poor conditions results in the slow decrease of viruses with high plasticity over time (Figure 6B). However, frequent periods of poor conditions lead to plastic viruses increasing in density over time (Figure 6C), despite having a lower lysis rate than invariant viruses over the entire duration of the simulation (due to an invariance cost of 0 in these simulations). We find that in environments with sufficiently short high-resource periods, plastic viruses increase in density over time when competing with invariant viruses (Figure S8). Notably, the fitness benefits of plasticity are dependent on a low non-viral death rate of susceptible and infected cells (Figure S8). If the cell death rate is too high, susceptible cell density quickly decreases during low-resource periods and both viruses go extinct. At intermediate death rates, all infected cells decrease in density during the no-mixing periods, reducing the advantage of plastic viruses. In essence, these simulations suggest that plasticity is favored in variable resource environments when very poor conditions are sufficiently common and plastic infected cells can survive them. Including a positive invariance cost would further favor high maturation rate plasticity across all resource conditions (Figure 5).

### Diversity in plasticity strategy promotes viral coexistence

Finally, we investigate whether variation in plasticity may allow for coexistence between viruses of different plasticity levels for non-zero invariance costs. Viruses with low plasticity can drive down host densities when resources are low, which increases resources and, in turn, benefits viruses with high plasticity. We identify coexistence through analysis of mutual invasion assuming that *ρ*_*r*_ = *ρ*_*η*_ (Figure 7A,B). We observe coexistence for a very limited part of the parameter space that results in large oscillations of virus densities. Within that parameter space, coexistence occurs for some pairs of *ρ*_*m*_ and *ρ*_*η*_ along the anti-diagonal (Figure 7C). In other words, complementary viral strategies are more likely to coexist than are trait-adjacent strategies, e.g., a highly plastic and a trait invariant virus may coexist whereas one of two highly plastic strategies would tend to dominate the other.

**Figure 7:**
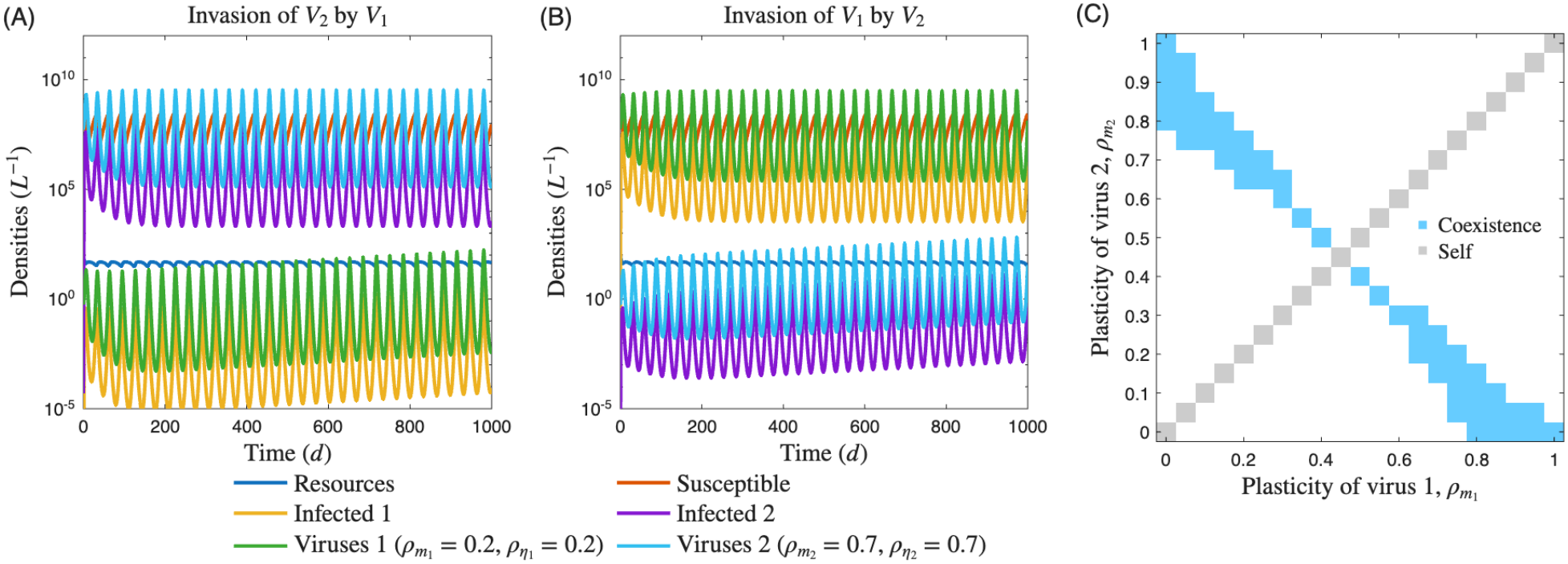
Coexistence between viruses with different plasticity. We determined coexistence by assessing whether two viruses of different plasticity could invade each other from an initial density of 1*L*^*−*1^. Due to large oscillations in viral dynamics, we defined invasion as an increase in maximum density over time (see Methods). Lysis rate plasticity *ρ*_*η*_ was set to maturation rate plasticity *ρ*_*m*_ for each virus. **(A)** Invasion of virus V2 (*ρ*_*m*_ = *ρ*_*η*_ = 0.7) by virus V1 (*ρ*_*m*_ = *ρ*_*η*_ = 0.2). **(B)** Invasion of virus V1 (*ρ*_*m*_ = *ρ*_*η*_ = 0.2) by virus V2 (*ρ*_*m*_ = *ρ*_*η*_ = 0.7). **(C)** Pairwise coexistence across maturation rate plasticity. Coexisting virus pairs are shown blue, with self-competition excluded in grey. Non-default parameter values are as follows: *S*_*out*_ = 10^6^ *L*^*−*1^, *R*_*out*_ = 50 *L*^*−*1^, *ω* = 0.5 *d*^*−*1^, *ϕ* = 5 × 10^*−*10^ *L* cell^*−*1^ *d*^*−*1^, *η*_*max*_ = 1.95 *d*^*−*1^, *m*_max,abs_ = 100 *d*^*−*1^, *δ*_*S*_ = 0.03 *d*^*−*1^, *λ* = 0.2.

## Discussion

Many viruses replicate in variable environments with resource-replete and resource-depleted periods. These changes in environmental resource availability impact host state, and in turn, the adaptive value of viral life history traits. Here, we developed an eco-evolutionary modeling framework that treats plasticity in viral maturation and viral lysis rates as continuous traits. We used this framework to estimate plasticity in experimental datasets of viruses of bacteria and phytoplankton and found high maturation rate plasticity and intermediate lysis rate plasticity across host-virus pairs. This consistent pattern motivated us to use an eco-evolutionary model to investigate phage, host and environmental conditions that drive the evolution of plasticity.

Our models revealed mechanisms by which plasticity can shape viral invasion fitness and competition across a range of environmental contexts. We found that increases in resource availability increased the degree of plasticity that maximizes viral fitness as measured through the basic reproduction number. In competition simulations, high resource influx also favored viruses with higher maturation rate plasticity. These findings are consistent with previous work showing an increase in phage performance for plastic viruses when resource input increases and cell growth rate decreases [14]. However, the optimal degree of plasticity is also shaped by the cost of trait invariance (i.e., the degree to which viruses can decouple replication traits from host metabolic state). For example, the maturation rate plasticity that maximize invasion fitness and growth increase with invariance cost.

From an ecological perspective, we identified host density and mixing rates (which control the turnover of populations and nutrients) as central in determining the lysis rate plasticity that maximizes viral invasion and growth. We observed that high lysis rate plasticity maximizes invasion fitness for a broad range of ecologically relevant host densities below a certain threshold. We would therefore expect high lysis rate plasticity in most natural settings, outside of extreme events like acute infections or blooms [28, 29, 30]. Viruses with high lysis rate plasticity also have advantages in variable environments and can survive nutrient poor conditions in their intracellular stage (i.e., within infected cells). High mixing rates increase cell turnover and therefore favor viruses that can complete an infection cycle quickly (short latent period), irrespective of nutrient conditions. This is consistent with existing work showing that the evolutionarily stable latent period decreases as the mixing rate increases (independently of maturation rate plasticity) [14].

While our findings clarify evolutionary drivers of plasticity, they raise questions about the underlying cellular mechanisms that generate maturation rate and lysis rate plasticity. The viral maturation rate depends on ribosome abundance, which in turn scales proportionally with growth rate. Cell size increases exponentially with cellular growth rate and could therefore drive a non-proportional scaling between cellular growth rate and viral maturation rate [31]. We observed this non-proportional scaling in a subset of experimental datasets [1, 3, 6, 10, 27]. Alternatively, differences in the elemental stoichiometric makeup of cells and viruses could also lead to non-linear relationships between cellular growth rates and viral maturation rates [32]. Many viruses also carry auxiliary metabolic genes – genes not essential to viral replication but that can modulate the host’s metabolism – to provide the resources needed for viral progeny [33], which would decrease maturation rate plasticity. Lysis rate, on the other hand, is governed by the molecular dynamics of proteins like holins [34]. Biophysical constraints may restrict both the minimum and maximum lysis rate away from its evolutionarily stable value and influence lysis rate plasticity.

Our work presents a flexible framework for incorporating plasticity and assessing its ecological and evolutionary consequences. However, the work also comes with caveats given the inclusion of multiple assumptions to simplify analysis. First, we limited our maturation rate plasticity parameter between 0 and 1, and assumed a linear relationship between maturation rate and cell growth rate, with a linear cost to invariance. In contrast, we estimated a small number of maturation rate plasticity values above 1 in our analysis of experimental data. This suggests a possible non-linear relationship between maturation rate and cell growth rate for certain virus-host pairs (see [14] for example). A non-linear relationship might also arise from the presence of an eclipse period, which is not incorporated in our model. Second, we surveyed a wide range of putative invariance costs. We are not aware of any quantitative measurement of the cost of trait invariance, which combines two processes: the decrease in maturation rate plasticity from an increase in viral genome size and the decrease in the maximum maturation rate from an increase in viral genome size. While maturation rates can be estimated from viral datasets [16], the degree of plasticity remains uncharacterized. Finally, we observed differences in plasticity values between limiting nutrients (e.g., especially vis a vis the impacts of iron on viral replication vs. host growth rates) that could motivate nutrient-specific mechanistic models.

Overall, this study explored the adaptive value of the degree of viral trait plasticity. In doing so, we found evidence for substantial maturation rate plasticity and lysis rate plasticity in experimental virus-microbe data. Our subsequent model analysis shows that both forms of plasticity should be favored across many environmental conditions, especially when cellular availability is low and/or variable. Moving forward, additional experimental work will be needed to assess the mechanistic basis for viral plasticity that, in turn, is likely to motivate new model developments that account for intracellular nutrient quotas and increased resolution of the viral life cycle. We anticipate that this growing body of experimental and modeling work will also help guide -omics based characterization of the extent and nature of viral plasticity across human, animal-associated, and environmental contexts.

## Acknowledgments

This work was funded by the Simons Foundation (SFI-LS-PROJECT-00011964) and supported by the University of Maryland’s Grand Challenges Grants program. J.S.W. and J.D.P are investigators at the University of Maryland-Institute for Health Computing, which is supported by funding from Montgomery County, Maryland, and The University of Maryland Strategic Partnership: MPowering the State, a formal collaboration between the University of Maryland, College Park, and the University of Maryland, Baltimore.

## Data availability

All code is available through github (https://github.com/jpourtois/viral_plasticity).

