## Supplemental Material for "Nutrient and host availability drive the evolution of viral plasticity"

### S1 Equilibrium analysis for virus-free model

We derived the analytical expression for the virus-free equilibrium $(R^*, S^*)$:

$$0 = \omega(R_{\text{out}} - R^*) - \epsilon \frac{\mu S^* R^*}{R^* + K} \tag{S1}$$

$$0 = \omega(S_{\text{out}} - S^*) + \frac{\mu S^* R^*}{R^* + K} - \delta_S S^* \tag{S2}$$

We first express $S^*$ in terms of $R^*$:

$$0 = \omega S_{\text{out}} - S^*(\omega + \delta_S) + \frac{\mu S^* R^*}{R^* + K} \tag{S3}$$

$$S^*(\omega + \delta_S - \frac{\mu S^* R^*}{R^* + K}) = \omega S_{\text{out}} \tag{S4}$$

$$S^* = \frac{\omega S_{\text{out}}}{(\omega + \delta_S - \frac{\mu S^* R^*}{R^* + K})} = \frac{\omega S_{\text{out}}(R^* + K)}{R^*(\omega + \delta_S - \mu) + \omega K + \delta_S K} \tag{S5}$$

The expression for $S^*$ is then used to obtain an expression for a quadratic function of $R^*$:

$$0 = \omega(R_{\text{out}} - R^*) - \epsilon \frac{\mu S^* R^*}{R^* + K} \tag{S6}$$

$$0 = \omega(R_{\text{out}} - R^*) - \epsilon \mu R^* \frac{\omega S_{\text{out}}}{R^*(\omega + \delta_S - \mu) + \omega K + \delta_S K} \tag{S7}$$

$$0 = (R_{\text{out}} - R^*)(R^*(\omega + \delta_S - \mu) + \omega K + \delta_S K) - \epsilon \mu R^* S_{\text{out}} \tag{S8}$$

$$0 = R^{*2}(\mu - \omega - \delta_S) + R^*(R_{\text{out}}(\omega + \delta_S - \mu) - K(\delta_S + \omega) - \epsilon \mu S_{\text{out}}) + R_{\text{out}} K(\omega + \delta_S) \tag{S9}$$

This can be solved for $R^*$ using the quadratic formula $R^* = \frac{-b \pm \sqrt{b^2 - 4ac}}{2a}$ where $a = (\mu - \omega - \delta_S)$, $b = R_{\text{out}}(\omega + \delta_S - \mu) - K(\delta_S + \omega) - \epsilon \mu S_{\text{out}}$ and $c = R_{\text{out}} K(\omega + \delta_S)$. These roots are always real for all real values of the parameters, i.e., $b^2 - 4ac > 0$:

$$\begin{aligned}
b^2 - 4ac &= [(\omega + \delta_S - \mu)R_{\text{out}} - K(\omega + \delta_S) - \epsilon\mu S_{\text{out}}]^2 - 4(\mu - \omega - \delta_S)(\omega + \delta_S)KR_{\text{out}} \\
&= (\omega + \delta_S - \mu)^2 R_{\text{out}}^2 + K^2(\omega + \delta_S)^2 + \mu^2\epsilon^2 S_{\text{out}}^2 \\
&+ 2R_{\text{out}}K(\omega + \delta_S)(\omega + \delta_S - \mu) + 2\mu\epsilon K S_{\text{out}}(\omega + \delta_S) - 2\mu\epsilon S_{\text{out}} R_{\text{out}}(\omega + \delta_S - \mu)
\end{aligned} \tag{S10}$$

Since $\mu$, $\epsilon$ $K$, $S_{\text{out}}$, $\omega$ and $\delta_S$ are all non-negative real numbers,

$$\begin{aligned}
b^2 - 4ac &= (\omega + \delta_S - \mu)^2 R_{\text{out}}^2 + K^2(\omega + \delta_S)^2 + \mu^2\epsilon^2 S_{\text{out}}^2 \\
&+ 2R_{\text{out}}K(\omega + \delta_S)(\omega + \delta_S - \mu) + 2\mu\epsilon K S_{\text{out}}(\omega + \delta_S) - 2\mu\epsilon S_{\text{out}} R_{\text{out}}(\omega + \delta_S - \mu) \\
&\geq (\omega + \delta_S - \mu)^2 R_{\text{out}}^2 + K^2(\omega + \delta_S)^2 + \mu^2\epsilon^2 S_{\text{out}}^2 \\
&+ 2R_{\text{out}}K(\omega + \delta_S)(\omega + \delta_S - \mu) - 2\mu\epsilon K S_{\text{out}}(\omega + \delta_S) - 2\mu\epsilon S_{\text{out}} R_{\text{out}}(\omega + \delta_S - \mu) \\
&= [(\omega + \delta_S - \mu)R_{\text{out}} + K(\omega + \delta_S) - \mu\epsilon S_{\text{out}}]^2
\end{aligned} \tag{S11}$$

So, $b^2 - 4ac \geq [(\omega + \delta_S - \mu)R_{\text{out}} + K(\omega + \delta_S) - \mu\epsilon S_{\text{out}}]^2 \geq 0$. The second inequality holds for all real values of the model parameters. Thus, both the roots of Equation S9 are real.

Further, at least one of these roots is positive: The sum of the roots of Equation S9 is $\Sigma = R_{\text{out}} + \frac{(\omega+\delta_S)K + \mu S_0}{\mu - \delta_S - \omega}$ and the product of the roots is $\Pi = \frac{\omega+\delta_S}{\mu-\delta_S-\omega} K R_{\text{out}}$. If $\mu > \delta_S + \omega$, then both $\Sigma$ and $\Pi$ are positive, implying that both roots of Equation S9 are positive. If $\mu < \delta_S + \omega$, then $\Pi$ is negative, which implies that exactly one root of Equation S9 is positive. In any case, atleast one of the roots of Equation S9 is positive.

The Jacobian, $J(R, S)$ for the virus-free system is given by:

$$J(R, S) = \begin{bmatrix} -\omega - \epsilon\mu S \frac{K}{(R+K)^2} & \frac{-\epsilon\mu R}{R+K} \\ -\frac{\mu S K}{(R+K)^2} & -\omega - \delta_S + \frac{\mu R}{R+K} \end{bmatrix} \tag{S12}$$

An equilibrium $(R^*, S^*)$ is stable if $\det(J(R^*, S^*)) > 0$ and $\operatorname{tr}(J(R^*, S^*)) < 0$.

$$\begin{aligned}
\operatorname{tr}(J(R^*, S^*)) &= -2\omega - \delta_S - \epsilon\mu S^* \frac{K}{(R^* + K)^2} + \frac{\mu R^*}{R^* + K} \\
&= -\omega - \omega\frac{S_{\text{out}}}{S^*} - \frac{\omega K}{R^*}\left(\frac{R_{\text{out}} - R^*}{R^* + K}\right)
\end{aligned} \tag{S13}$$

$$\det(J(R^*, S^*)) = \tag{S14}$$

For any equilibrium $(R^* > 0, S^* > 0)$, we can show that $R^* < R_{\text{out}}$ using Equation S1. Then $\operatorname{tr}(J(R^*, S^*)) < 0$.

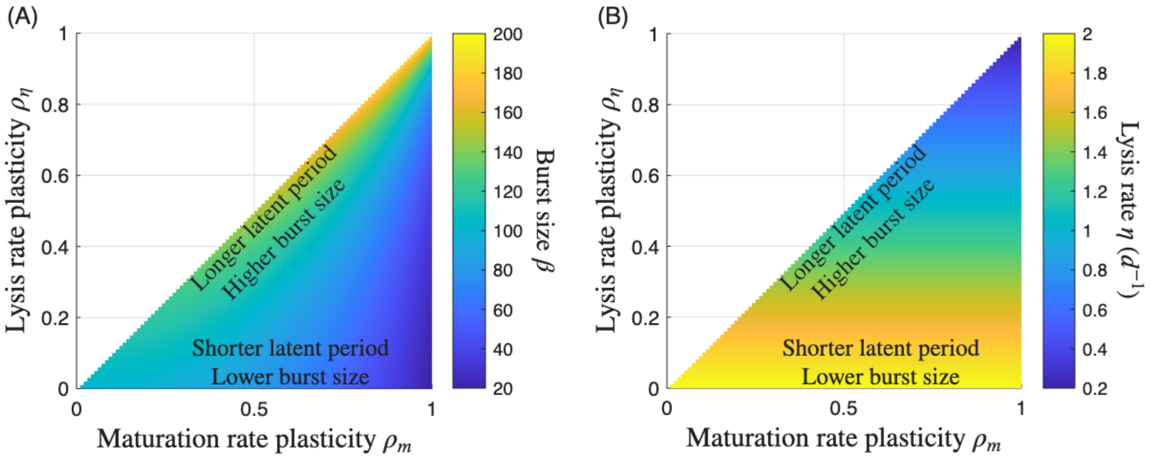


**Figure S1: Model description**. Tradeoff in burst size $\beta$ **(A)** and lysis rate $\eta$ **(B)** arising from maturation rate plasticity $\rho_m$ and lysis rate plasticity $\rho_\eta$ for invariance cost $\lambda$ of 0.5. We limit our analysis to lysis rate plasticity arising from maturation rate plasticity and do not consider the parameter space $\rho_\eta > \rho_m$.

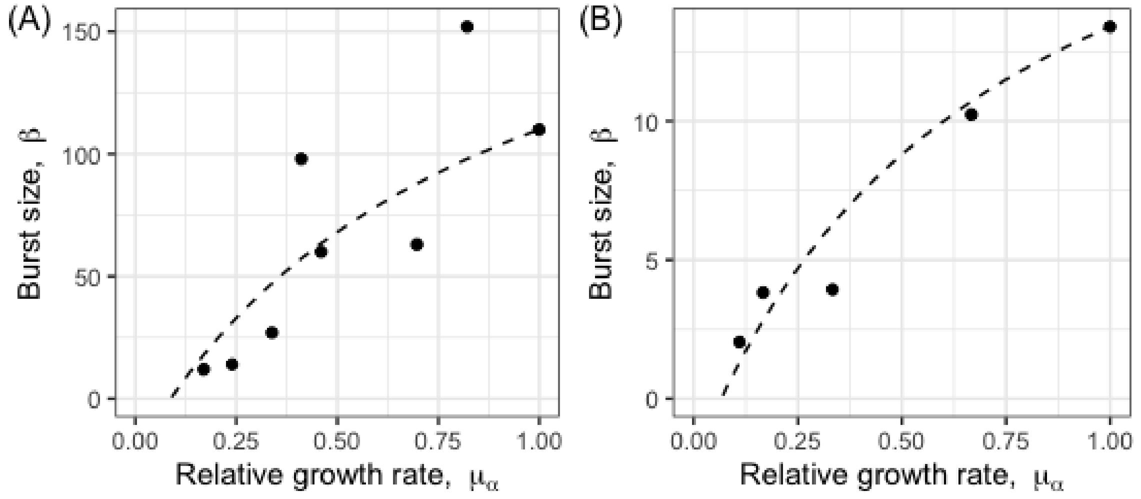


**Figure S2: Observed and predicted burst size across relative bacterial growth rate for phage T4 and *E. coli*.** **(A)** Burst size of phage T4 infecting *E. coli* in different media [1] **(B)** Burst size of phage T4 infecting *E. coli* in phosphate-buffered minimal media with glucose at different dilution rates [2]. Growth rates were normalized with respect to the maximum growth rate for each experiment. Predicted burst size (dotted line) was calculated from predicted maturation rate and lysis rate based on estimated plasticity $\rho_m$ and $\rho_\eta$.

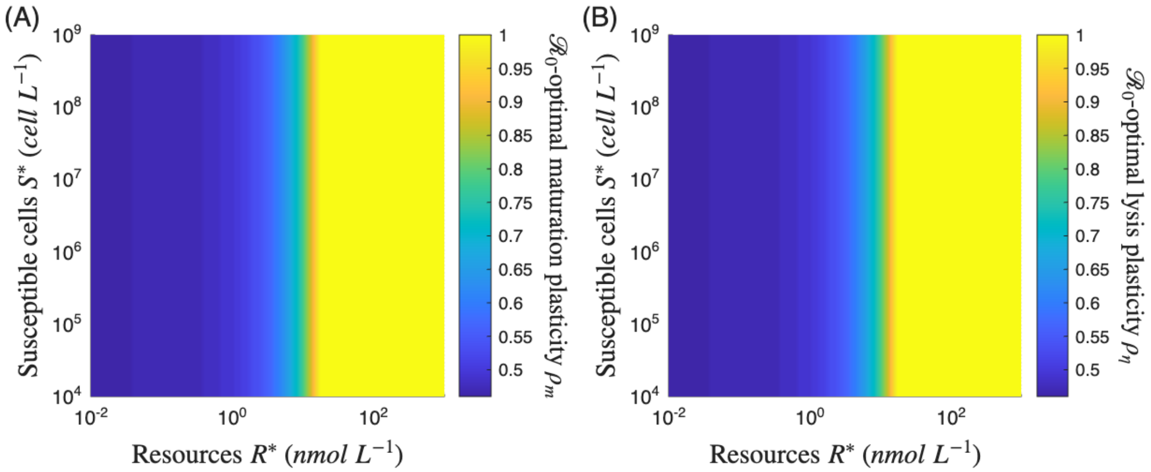


**Figure S3: $\mathcal{R}_0$-maximizing maturation rate and lysis rate plasticity**. Pair of maturation rate plasticity $\rho_m$ **(A)** and lysis rate plasticity $\rho_\eta$ **(B)** that maximize $\mathcal{R}_0$ for invariance cost $\lambda = 0.5$.

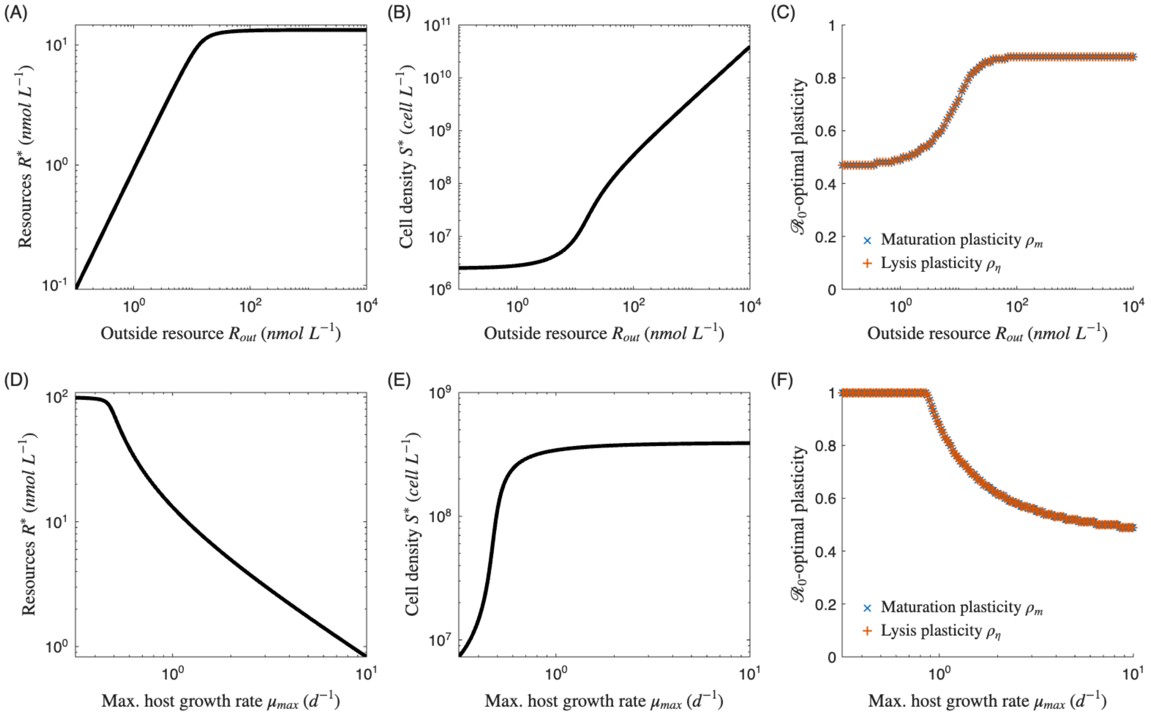


**Figure S4: $\mathcal{R}_0$-maximizing plasticity across outside resource concentration $R_{out}$ and maximum host growth rate $\mu_{max}$.** **(A)** Equilibrium resource concentration $R^*$ across outside resource concentration $R_{out}$. **(B)** Equilibrium susceptible host density $S^*$ across outside resource concentration $R_{out}$. **(C)** Maturation rate plasticity $\rho_m$ and lysis rate plasticity $\rho_\eta$ that maximize $\mathcal{R}_0$ at equilibrium across outside resource concentration $R_{out}$. **(D)** Equilibrium resource concentration $R^*$ across maximum host growth rate $\mu_{max}$. **(E)** Equilibrium susceptible host density $S^*$ across maximum host growth rate $\mu_{max}$. **(F)** Maturation rate plasticity $\rho_m$ and lysis rate plasticity $\rho_\eta$ that maximize $\mathcal{R}_0$ at equilibrium across maximum host growth rate $\mu_{max}$. Invariance cost $\lambda$ is set to 0.5 for all panels.

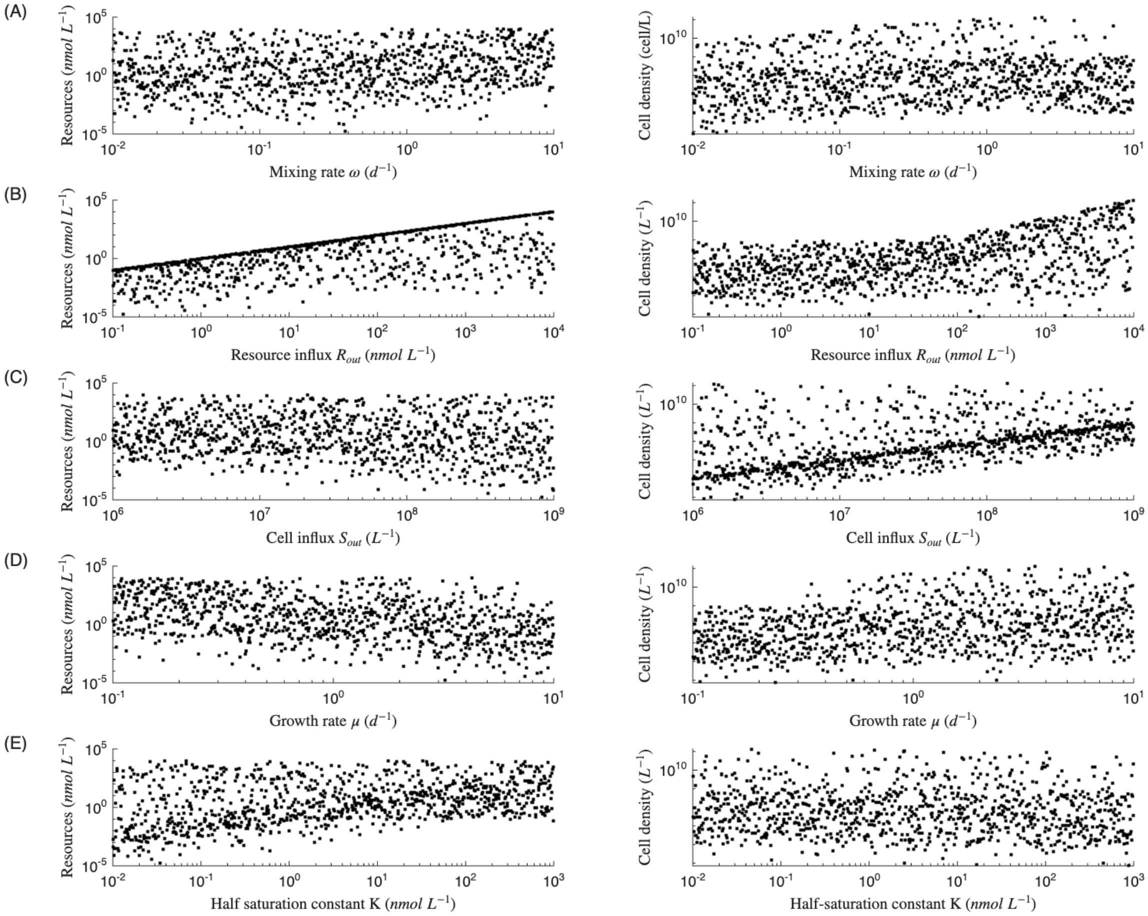


**Figure S5: Sensitivity analysis for virus-free model.** We generated parameter sets through Latin Hypercube Sampling across ranges shown here. Equilibrium values for resources (left) and susceptible cell densities (right) were determined analytically (see Methods). We calculated Pearson correlation coefficients between each parameter and resources concentration ($r_R$) or susceptible cell density ($r_S$).**(A)** Mixing rate $\omega$ ($r_R = 0.15$, $r_S = -0.030$). **(B)** Resource influx $R_{out}$ ($r_R = 0.65$, $r_S = 0.56$). **(C)** Susceptible cell influx $S_{out}$ ($r_R = 0.025$, $r_S = -0.024$). **(D)** Maximum cell growth rate $\mu_{max}$ ($r_R = -0.16$, $r_S = 0.13$). **(E)** Half saturation constant $K$ ($r_R = 0.063$, $r_S = -0.040$).

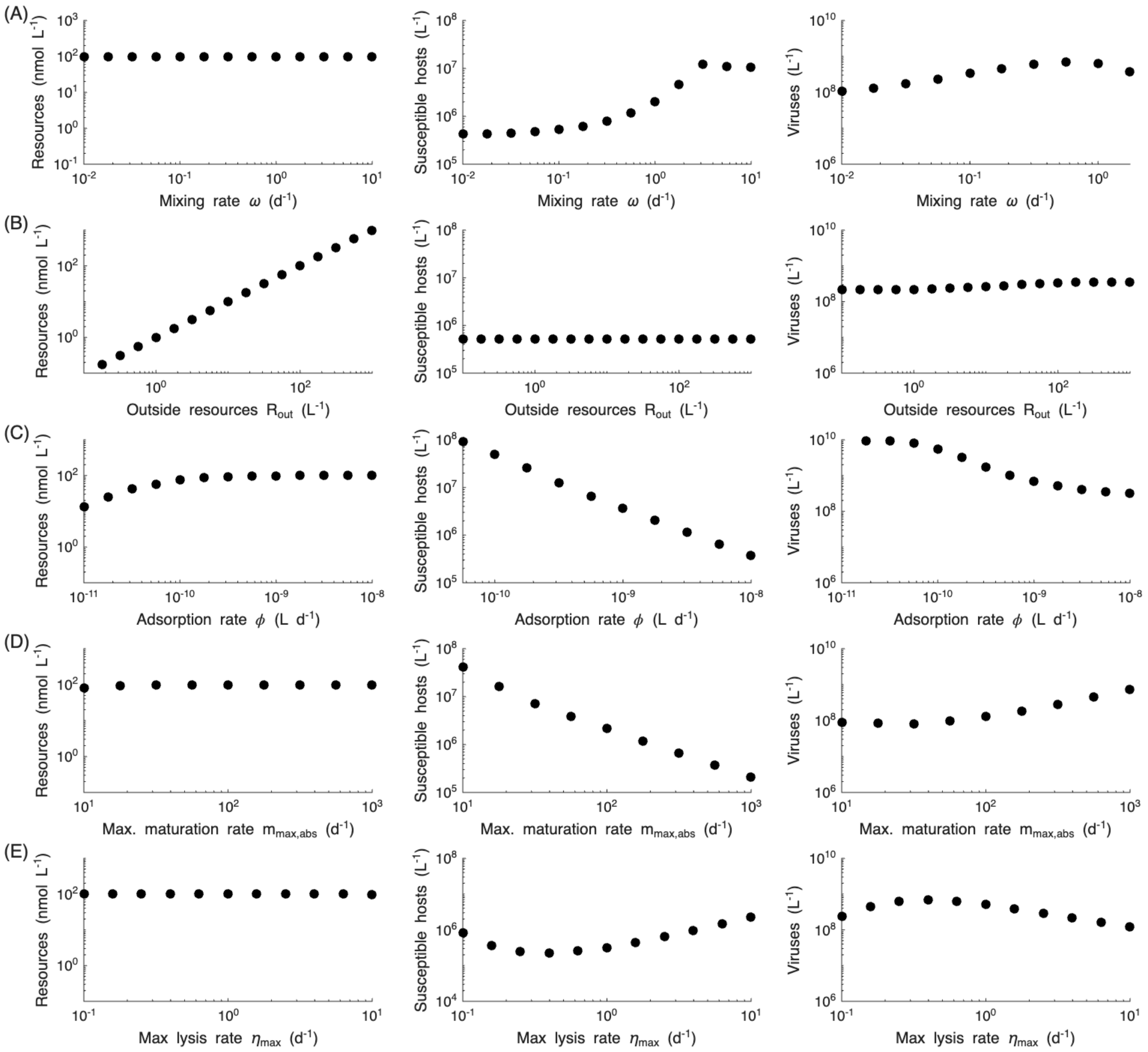


**Figure S6: Sensitivity analysis for model with viruses.** Equilibrium values for resources (left), susceptible hosts (middle) and viruses (right) were obtained by averaging the last 400 days of 1000-day simulations. Parameters were varied one at a time while keeping all others fixed (see default values in Table 1). **(A)** Mixing rate $\omega$. **(B)** Resource influx $R_{out}$. **(C)** Adsorption rate $\phi$. **(D)** Maximum maturation rate $m_{max,abs}$. **(E)** Maximum lysis rate $\eta_{max}$.

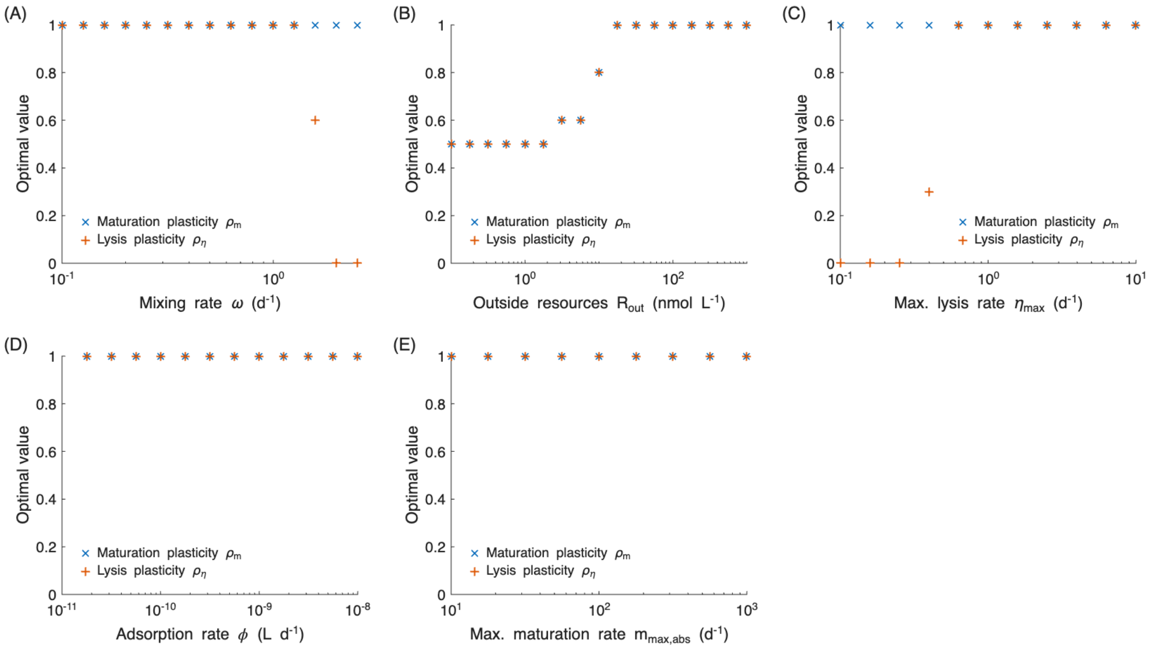


**Figure S7: Competition simulations in chemostat with constant resource and susceptible cell inflow.** Maturation rate plasticity $\rho_m$ and lysis rate plasticity $\rho_\eta$ of the virus with the highest density at the end of a 3000-day competition simulation are shown. All initial virus density were set to $10^7 L^{-1}$ and were introduced in a virus-free system at equilibrium. Invariance cost is set to 0.5. **(A)** Mixing rate $\omega$. **(B)** Outside resource concentration $R_{\text{out}}$. **(C)** Maximum lysis rate $\eta_{\text{max}}$. **(D)** Adsorption rate $\phi$. **(E)** Maximum maturation rate $m_{\text{max,abs}}$.

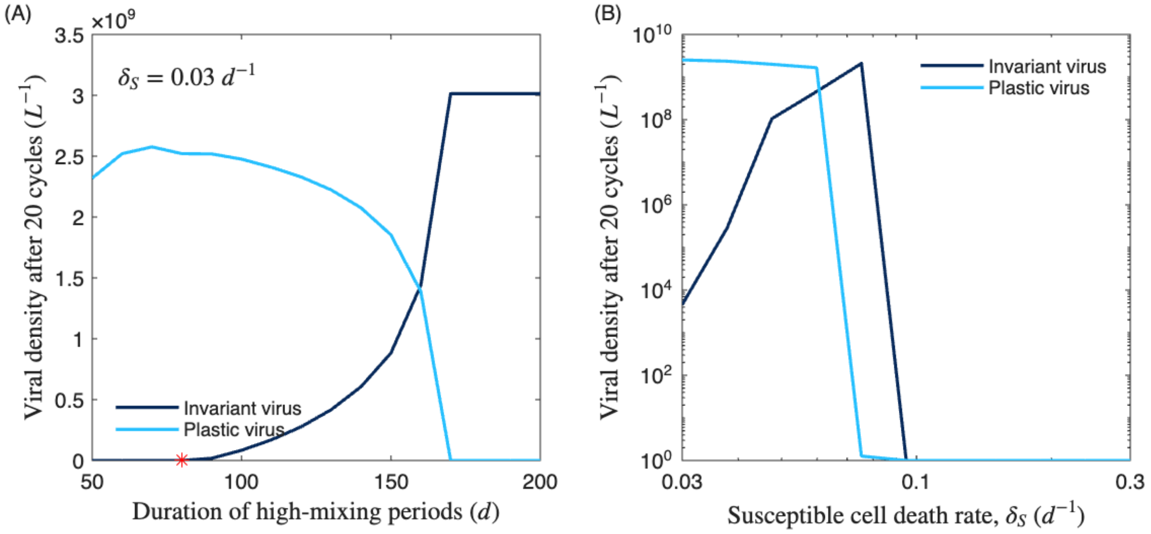


**Figure S8: Effect of high-mixing period duration and host death rate on fitness of plasticity**. **(A)** Density of plastic and invariant viruses after 20 cycles of low/high resource cycles across a range of duration of high-resource periods, with low-resource period duration fixed at 50 days. Red star shows duration used in the right panel. **(B)** Density of plastic and invariant viruses after 20 cycles of low/high resource cycles for a range of susceptible cell death rates, with high-resource and low-resource periods lasting 80 and 50 days respectively.
